# Exploratory study of pulsed electric field ablation on atherosclerotic plaque in a rabbit model

**DOI:** 10.1101/2023.12.05.570315

**Authors:** Ye Xuying, Hu Jiashen, Cao Shisheng, Xu Xinyu, Xue Zhixiao, Lu Chengzhi, Yin Huijuan

**Affiliations:** Department of cardiology, Tianjin First Central Hospital, Tianjin, 300192; Institute of Biomedical Engineering, Chinese Academy of Medical Sciences & Peking Union Medical College, Tianjin, 300192; School of Biomedical Engineering and Technology, Tianjin Medical University, Tianjin, 300072

**Keywords:** Pulsed field ablation, Atherosclerosis, Hyperlipidemia, Vascular remodeling, Cardiovascular disease

## Abstract

New understanding of the pathogenesis of atherosclerotic diseases has led to the emergence of new therapeutic approaches. We explored the potential therapeutic effects of pulsed field potential ablation (PFA), a non-thermal ablation technique with high tissue selectivity, on atherosclerotic plaques. Carotid arteries of 30 high-fat rabbits were dilated with a balloon to obtain atherosclerotic plaques. PFA was administered on the carotid atherosclerotic plaques with 1000V/cm, 2000V/cm, and 1000V/cm ablation followed by rapamycin infusion. There were no visible changes in blood vessels after acute ablation, but apoptosis and polarity of cells were observed in atherosclerotic plaques. At 7 and 30 days after ablation, the density of lipid deposition in the plaque was significantly reduced, and multiple layers of new arranged anterograde smooth muscle cells appeared, replacing the original atherosclerotic plaque. The residual atherosclerotic structure is sandwiched between the new smooth muscle layer and the original smooth muscle layer, which makes vascular wall thicker and makes vascular wall elasticity increased. Rapamycin delays the vascular remodeling process. Conclusion: PFA ablation can reduce lipid deposition in atherosclerotic plaques, cause vascular remodeling, and enhance vascular elasticity. We believe that it may be a potential method for the treatment of atherosclerotic plaques.

## 1. Introduction

Atherosclerotic diseases, including coronary heart disease, stroke, and peripheral vascular disease, etc., are the leading killers of death worldwide. According to data reported by the American Heart Association in 2023, the prevalence of cardiovascular disease (CVD) (comprising coronary heart disease, heart failure, stroke, and hypertension) in adults ≥20 years of age was 48.6% overall (127.9 million in 2020) and increases with age in both males and females^[1]^. With the discovery of new evidence on the pathology and pathogenesis of atherosclerosis, atherosclerosis(AS) is no longer considered an inevitable development of aging, but a reversible and manageable disease^[2]^. Superficial erosion is considered as an increasing cause of arterial thrombosis^[3]^, thus becoming a new target of atherosclerosis.

Pulsed electric field ablation (PFA) is a new ablation method that has been developed in recent years and has been used in the treatment of tumor ablation ^[4,5]^and atrial fibrillation ablation^[6]^. PFA uses high-voltage electrical pulses released between electrodes to form irreversible electroporation on the cell membrane, increasing cell permeability and leading cell to death in the end^[7]^. Most importance of all, due to the difference of resistance of biological tissues, PFA has remarkable tissue specificity. Our previous studies have shown that cellular tissues such as tumors, cardiomyocytes and smooth muscle cells are more sensitive to PFA than connective tissues^[8, 9]^. Compared with normal arteries, atherosclerotic arteries have different structure and lower resistance characteristics. So, in this study, we performed PFA on atherosclerotic carotid arteries in rabbits to explore the possibility of PFA ablation for the treatment of atherosclerotic arteries.

## 2. Methods and materials

### 2.1 Materials

The high frequency alternating pulsed field ablation equipment and electrode device were supported by Tianjin Intelligent Health Medical Co., Ltd.

24G disposable intravenous indenture needles were purchased from Guangdong Baihe Medical Technology Co.,Ltd. The coronary guided wire needles were purchased from Meritmedical Systems,Inc, America. SION BLUE coronary guide wires were purchased from Asahi Intecc Co., Ltd. The 3.0*20 balloons were purchased from Boston Scientific Corporation.

Sumianxin (Xylazine Hydrochloride Injection, 2ml:0.2g) was purchased from Changsha Best Biological technology institute CO., LTD, and Urethane (S11036-500g) was purchased from Shanghai yuanye Bio-Technology Co., Ltd.

The primary antibodies PCNA antibody and Apolipoprotein A1/APOA1 antibody (2G4)(sc-69755), were purchased from Santa Cruz Biotechnology, Inc., USA, α-SMA antibody (Ab7817) was purchased from Abcam, and PCNA antibody was purchased from Bosterbio. The second antibody MaxVision™ HRP-Polymer anti-Mouse HIC kit (KIT-5002) was purchased from MXB Blotechnologies, China. TUNLE kit (KGA702) was purchased from KeyGEN BioTECH, China. HE staining reagents (YM1001) purchased from Nanjing Yorm Biotechnology Co., Ltd, China, and EVG staining reagents (G1596) from Beijing Solarbio Science & Technology Co., Ltd., China.

#### 2. Animal model

Thirty Japanese white rabbits (average 2.5kg weight, half male and half female) were purchased from Beijing HFK Bioscience CO., LTD. All experimental protocols with all rabbits were approved by the animal ethics and welfare committee (approval number: 2022241) of Institute of Radiological Medicine, Chinese Academy of Medical Sciences. The rabbits were fed with 100g high fat food (formula: 5% lard, 1% cholesterol, 0.2% sodium cholate, 5% egg yolk powder, 88.8% base feed) per day for 3 months to induce hyperlipidemia. Four items of blood lipid including TG, TC, HDL and LDL were detected before (F0), 1 month (F1M) and 3 months (F3M) after high fat feeding. After a month of high fat feeding, balloon dilatation was performed on the left or right carotid artery of the rabbits under X-ray guidance. In detail, the rabbit was anesthetized (0.25mL/kg·weight Sumianxin || by intramuscular injection followed by 30% Urethane *i*.*v*.) and immobilized head-on to the operating table of Digital subtraction angiography machine. 24G disposable intravenous indenture needle was punctured into rabbit middle ear artery (left or right ear) to establish arterial passage. The coronary guided wire needle was inserted into the tail of the indenture needle, and then the SION BLUE coronary guide wire was introduced into the carotid artery through the passage. The 3.0*20 balloon was sent into the carotid artery along the guide wire, and the intended dilatation site was determined by angiography (between 4 to 6 cervical vertebrae). Balloon dilatation was performed as follows: filling the balloon 60s with 10ATM and shrinking the balloon 60s, repeated three times. Three consecutive parts were dilatated, and the total length up to 6cm. The rabbits continued to be fed high fat after surgery.

#### 3. Vascular ultrasound

After 3 months of high-fat feeding, and 2 months after balloon dilation, rabbits underwent vascular ultrasonography under muscular anesthesia. Rabbit neck was shaved and fully exposed, and both left and right carotid arteries were subjected to vascular ultrasound. Record the diameter of carotid artery, and observe the intima and wall of blood vessels to determine whether there are atherosclerotic plaque features.

#### 4. Pulsed electric field ablation

Rabbits with successful carotid atherosclerotic plaque modeling were randomly divided into 3 groups with 9 rabbits in each group. Pulsed electric field ablation (PFA) was performed on the carotid artery suffered balloon dilation. The rabbits were anesthetized, the skin and muscle layers of the neck were cut open, and the carotid artery was carefully separated. The flexible electrodes surround the blood vessel, and the distance between the positive and negative electrodes is 1cm. 1000V and 2000V voltages were performed to different groups of rabbits, called PFA1000V, PFA2000V and PFA1000V+RAPA groups. Pulsed electric field ablation works in a bi-directional short pulse mode, with 5 microseconds for each positive and negative pulse. A total of 100 pulses are implemented, and the average working current is 0.8A. In PFA1000V+RAPA group, the carotid artery was clipped at both ends after 1000V ablation and 100 *μ*L 0.15mg/ml rapamycin was injected with a 2.5mm needle syringe. Blood flow is restored in 90 seconds. After ablation, the wound was sutured. High fat feeding continued after operation.

#### 5. Histological analysis

At three time points immediately, on the 7th and 30th days after PFA operation, 3 rabbits in each group were sacrificed, and the left and right carotid arteries were quickly obtained, fixed with 4% paraformaldehyde, embedded in paraffin, and sliced. HE staining, TUNEL staining and EVG staining were used to detect the necrosis, apoptosis and elastic fiber changes of carotid artery induced by PFA, respectively. Immunohistochemical staining was performed with Apolipoprotein A I/APOA1 antibodies (2G4) and α-SMA to detect lipids in atherosclerotic plaques according to the manufacturer’s protocol.

The image acquisition was obtained by the digital pathology scanner (NanoZoomer S60, Hamamatsu, Japan) under 20× magnification.

#### 6. Statistical analysis

All data were presented as the mean ± standard deviation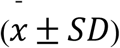. Comparisons between groups were performed using one-way ANOVA. Significance was set at p<0.05. Statistical analyses were performed using Origin2021.

## 3. Results

### 3.1 PFA performed on carotid atherosclerosis

The work flow and balloon dilatation were shown as Figure 1A and 1B. After 1 month of high-fat diet feeding, interventional balloon dilatation was performed through the rabbit middle ear artery to cause dilatation injury of the carotid artery, and then high-fat feeding was continued until 3 months. Vascular ultrasound was used to verify carotid artery dilation. The results showed that carotid artery diameter was obviously widened after balloon dilation. Compared with contralateral carotid arteries, 80 percent of the 30 rabbits had dilated arteries, and 35 percent had dilated arteries greater than 1.2 times (Figure 1C). After surgical exposure of the carotid artery, we observed that the dilated carotid artery was not only thick in diameter ratio to the side, but also white and thickened in wall with greasy feeling. These features all indicate the establishment of a carotid atherosclerosis model. We also tested four lipid indicators, including triglyceride (TG), total cholesterol (TC), low density lipoprotein (LDL) and high density lipoprotein (HDL). The results showed that after high fat feeding, these four indexes were significantly increased, in which the increase of LDL and TC was more than 10 times (Figure 1D). These serological indicators of hyperlipidemia provide basic conditions for the formation of carotid atherosclerotic plaques in rabbits. PFA was performed on the side of the carotid artery suffered balloon dilatation surgery in the rabbits within 1 week after ultrasound. (Figure 1E).

**Figure 1.**
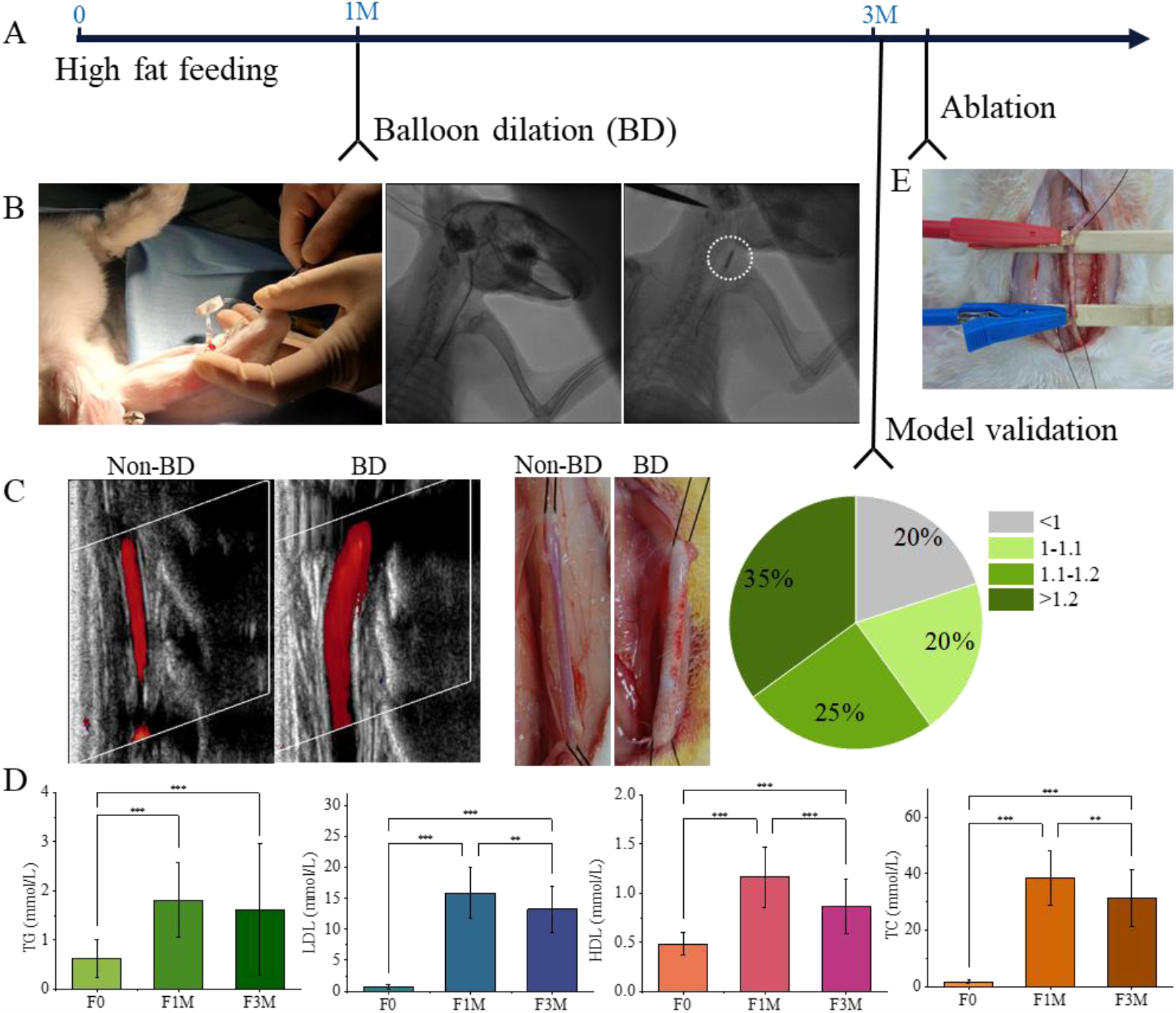
Establishment and validation of carotid atherosclerotic plaque model in rabbits. A. The flow scheme of the rabbit model building and Pulsed electric field ablation. The rabbits were fed with high fat through the whole process. B. Balloon dilation was performed on the left or right carotid artery through rabbit central ear artery under X-ray guidance after one-month high fat feeding. C. Vascular ultrasonography was used to validate the carotid dilatation as the performance of atherosclerosis. Images of exposed blood vessels also demonstrated atherosclerosis. Non-BD means no treatment of balloon dilation and BD means treatment of balloon dilation. The pie chart represents the ratio of BD molded vessels to the diametrically untreated vessels on the opposite side. D. The four indexes of blood lipid showed significant increase before high fat feeding (F0) and at 1 and 3 months (F1M and F3M) after feeding. Significant difference: *p<=0.05, **p<=0.01, ***p<=0.001. E. Pulsed electric field ablation was performed within 1 week after confirmation of the carotid atherosclerosis model. The distance between the two electrodes is 1cm.

### 3.2 Blood vessel morphology changes after PFA ablation

In the surgical field of view, we did not find significant structural changes in blood vessels within half an hour after ablation, except that some blood vessels became redder in color. (Figure 2, post-PFA 0.5h column). However, the blood vessels significantly thickened and their elasticity weakened after 7 days of PFA treatment, (Figure 2, post-PFA 7d column). After 30 days, the blood vessels still look thick, shiny and white, and elastic to the touch. (Figure 2, post-PFA 30d column). HE staining was performed on these vascular tissues and it was found that at all observation time points, the vascular lumen remained unobstructed and no significant stenosis or obstruction was observed (Figure 2, embedded images).

**Figure 2.**
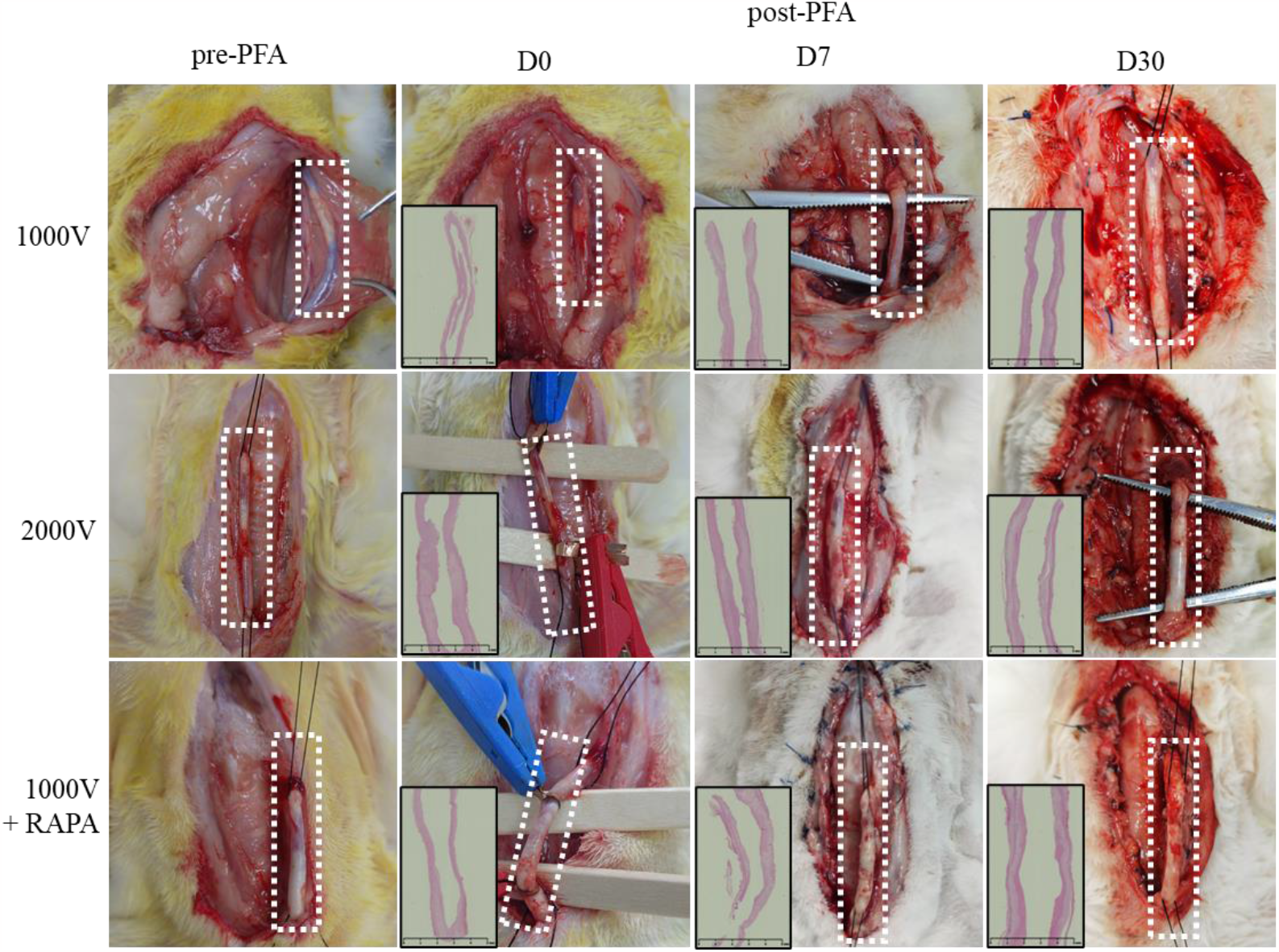
The change of blood vessel morphology after PFA ablation in different stages. The white dotted box shows the vessels being ablated. The embedded image was HE staining of the longitudinal section of the vessel.

In order to observe the effect of ablation on the vascular microstructure, we performed a variety of histological staining on the vessels half an hour after ablation, including HE staining, EVG staining, TUNEL staining and α-SMA immunohistochemical staining. As shown in Figure 3, we first compared the effect of 2000V ablation on normal and atherosclerotic vessels. After 2000V ablation of normal blood vessels, basically no structural changes occurred, the endothelia remained intact, smooth muscle was arranged along the vascular axis, and the elastic fiber structure maintained continuity and density. In contrast, the vascular morphology of atherosclerotic vessels changed significantly after ablation at 2000V. There was obvious apoptosis in the plaque tissue of the inner cortex, and the smooth muscle cells and elastic fibers showed obvious polar arrangement towards the vascular axis. The smooth muscle cells in the medium layer also showed apoptosis, and the cell structure was destroyed, but the elastic fibers still maintained the original sparse structure caused by lipid deposition. The vascular structure is obviously dehydrated, so that tissue rupture occurs during the preparation of tissue sections. Compared with 2000V ablation, 1000V ablation (including the addition of rapamycin) resulted in more minor changes in vascular structure, only the polar arrangement of smooth muscle cells and elastic fibers in the plaque could be seen, but no apoptosis of smooth muscle cells was observed.

**Figure 3.**
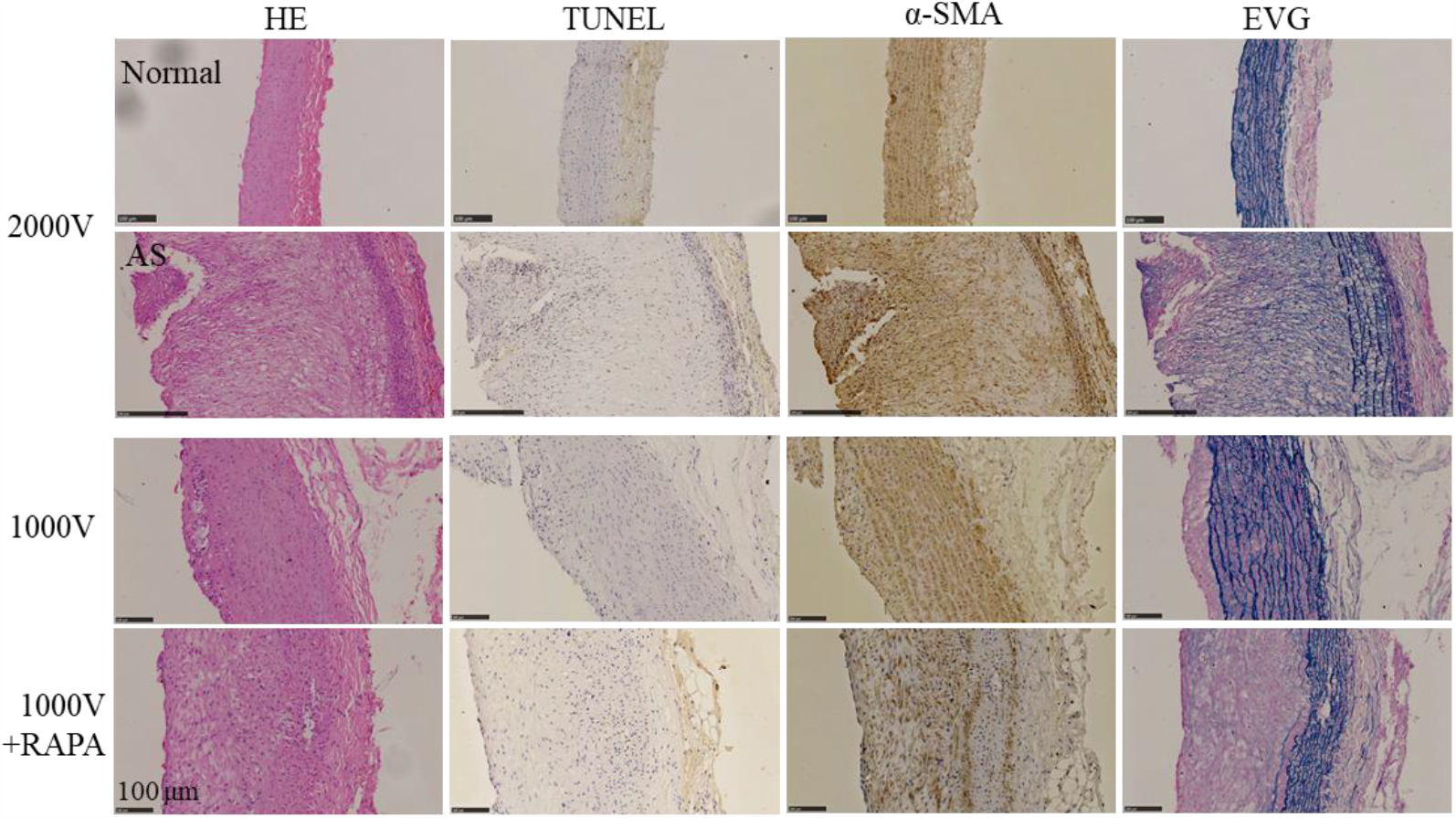
Vascular pathological changes at half an hour after PFA ablation. HE and TUNEL were used to observe the destruction of cells in all layers of blood vessels, SMA marked the actin of smooth muscle cells, EVG marked the elastic fibers.

### 3.3 Inhibition of lipid deposition by PFA

Lipid deposition is a key indicator of atherosclerotic plaque, and we observed the inhibitory effect of PFA ablation on vascular lipid deposition. As shown in Figure 4, all three ablation regimens of PFA saw a significant reduction in lipid deposition after 30 days. The reduction of lipid deposition does not occur immediately but is a long process. We could still see some lipid deposition in the blood vessels until the 7th day after PFA ablation. By day 30, however, l Lipid deposition decreased significantly. In addition, we also observed that rapamycin enhanced the inhibitory effect of PFA ablation on lipid deposition. Compared with the CP group, lipid deposition decreased by 28.9% in the 2000V ablation group and 22.6% in the 1000V ablation group. Most importantly, lipid plaques disappeared after PFA treatment.

**Figure 4.**
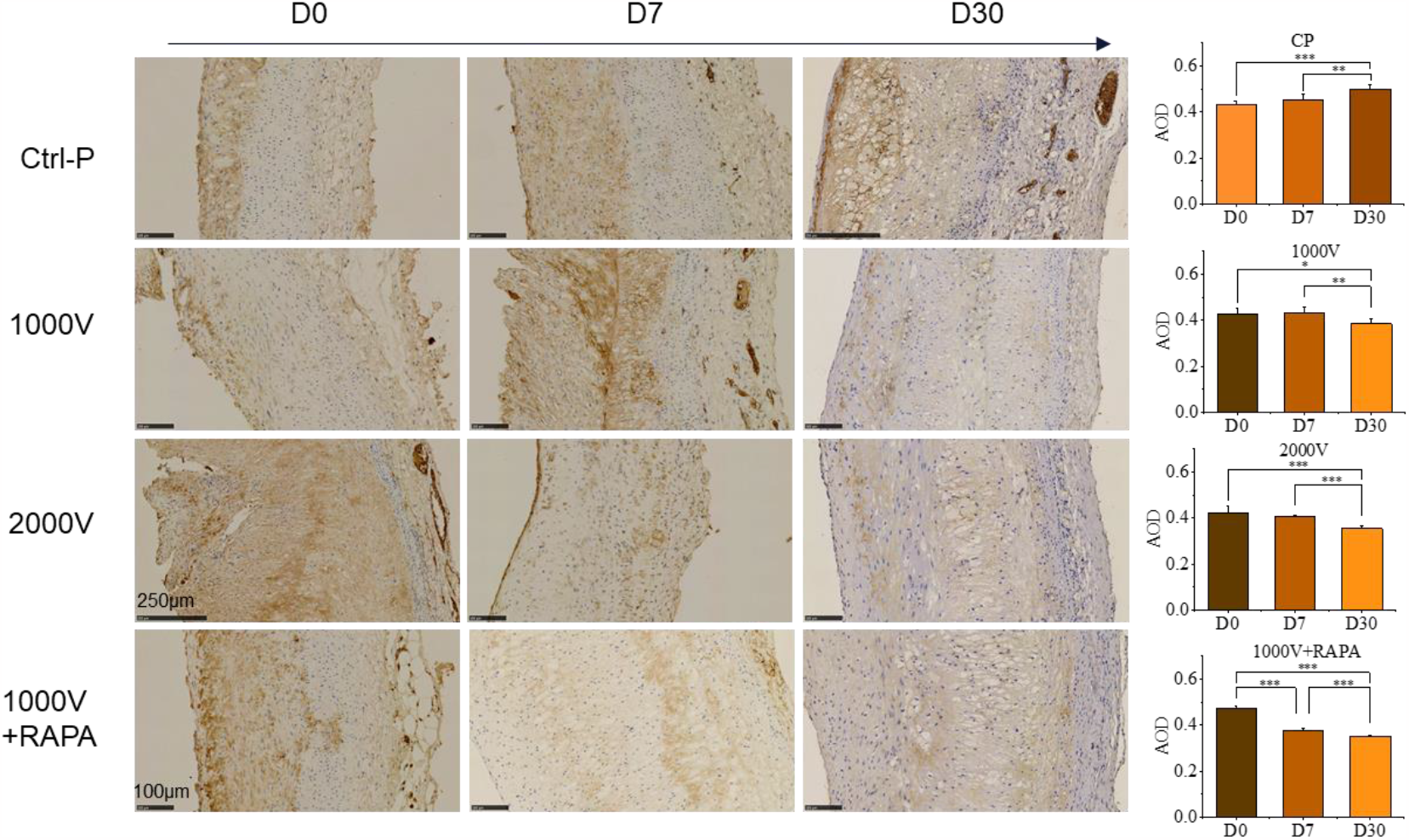
Inhibition of lipid deposition by PFA ablation. The rows represent the different groups and the columns represent the time after ablation. The average optical density (AOD) of lipid deposition was quantitatively described by the histogram. Significant difference: ^*^p<=0.05, ^**^p<=0.01, ^***^p<=0.001.

### 3.4 Vascular remodeling after PFA

What intrigued us was the reason for the decrease in lipid deposition. We first observed the active proliferation of cells in the vascular tissue after ablation, which may be the cause of lipid elimination. As shown in Figure 5, we found that the proliferative activity of cells varied significantly with different PFA treatments. Half an hour after the ablation, there was a significant increase in active proliferating cells in the 2000V ablation group. We considered that these are inflammatory cells recruited in response to vascular damage caused by 2000V ablation. 1000V ablation showed a peak of proliferating cells on day 7. It is considered that these are newborn smooth muscle cells, which is confirmed by Figure 6. Rapamycin delayed the proliferation of such nascent smooth muscle cells, which were observed to prolifically differ from those in the PFA group at 30 days after ablation.

**Figure 5.**
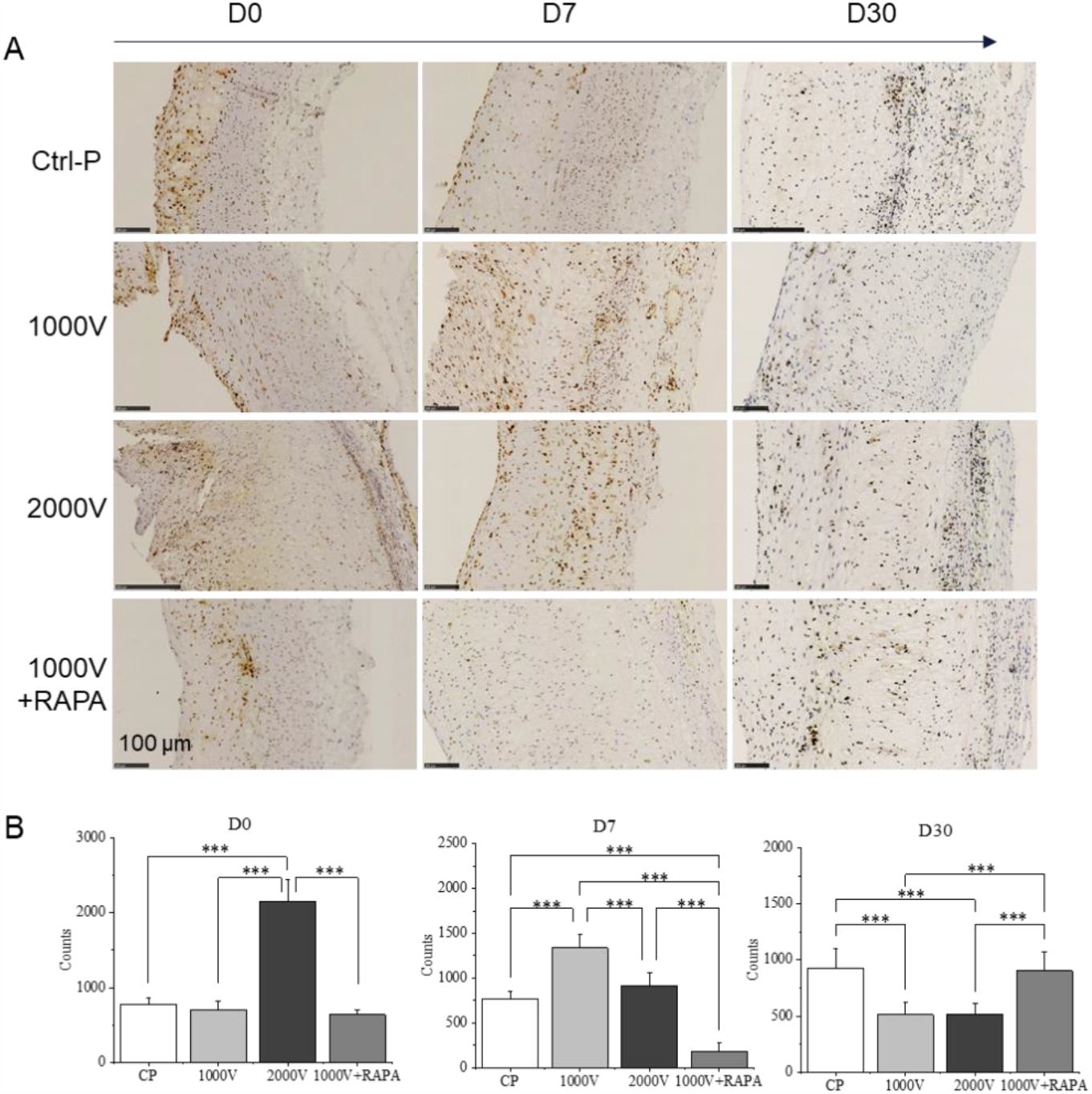
The proliferative cells in blood vessels after PFA ablation labeled by PCNA. A. PCNA stained HIC images of each group; B. Number of PCNA positive cells in vessels of the same length. Significant difference: ^*^p<=0.05, ^**^p<=0.01, ^***^p<=0.001.

**Figure 6.**
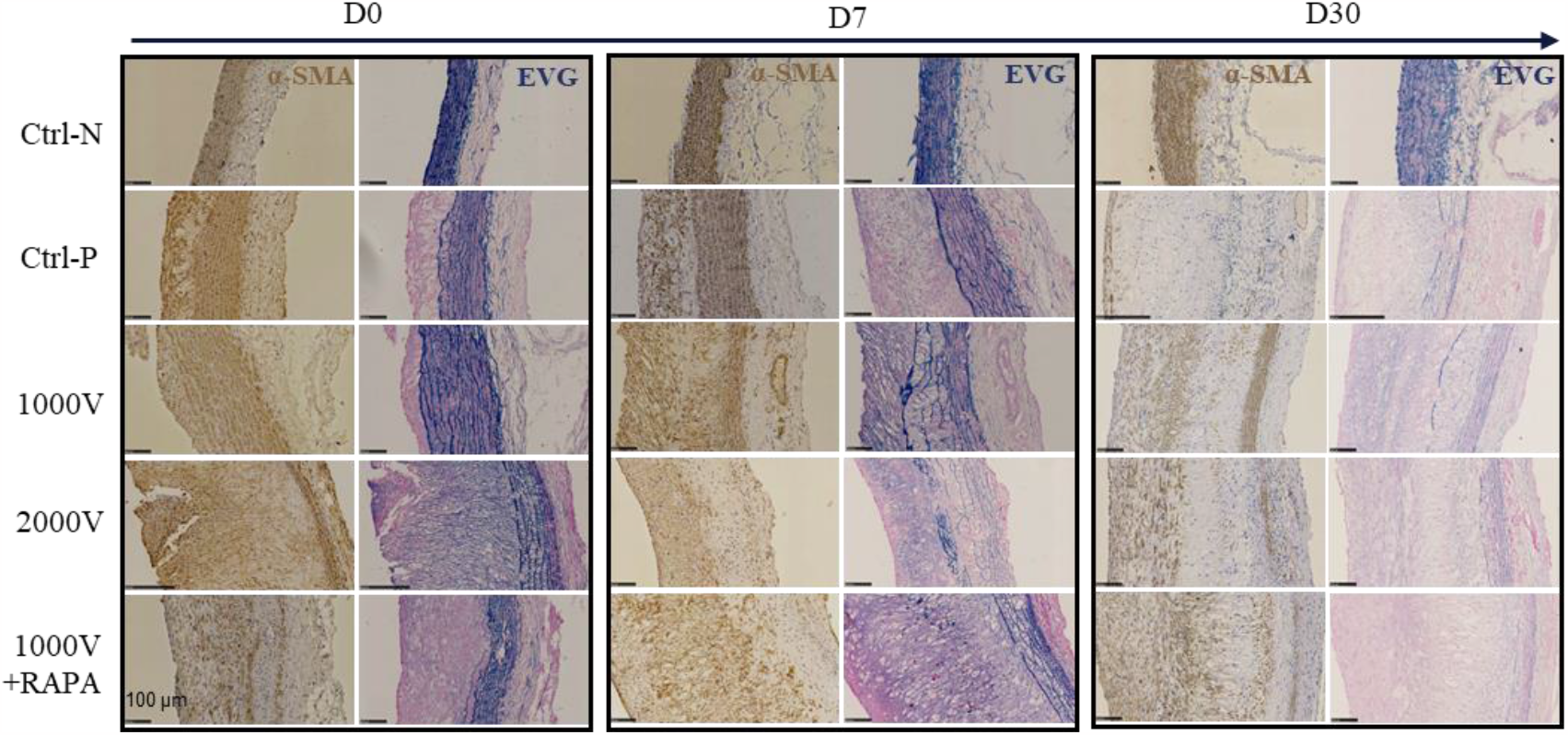
The process of vascular remodeling after ablation. HIC images with α-SMA marking smooth muscle cells and EVG staining marking elastic fibers (blue).

Inflammatory cells and new smooth muscle cells may be responsible for the decrease in lipid deposition. At the same time, the new smooth muscle cells reshaped the vascular structure. Figure 6 compared the changes of smooth muscle cells and elastic fibers in each group at 30 days after ablation. The most significant process of vascular remodeling occurred in 2000V. Immediately after ablation, the elastic fibers at the plaque site were aligned in the radial direction of blood vessels under the influence of electric field. On day 7 after ablation, the new smooth muscle and elastic fibers were clearly arranged along the radial direction of the blood vessels. By day 30, the new multilayer smooth muscle structure was basically remodeled. There is loose connective tissue between the new smooth muscle structure and the remaining original smooth muscle structure, and we estimate that the complete remodeling of blood vessels may take longer. Vessels ablated at 1000V undergo a similar but slow remodeling process, with rapamycin making the whole process even slower.

## 4. Discussion

Our previous studies on PFA for atrial fibrillation have shown that PFA ablation has a selective difference between cardiomyocytes and smooth muscle cells, with the latter more tolerant to PFA than the former^[9]^. We also proved that the same dose of PFA ablation would not cause damage to the surrounding connective tissue and nerve tissue. Other studies have also reached a similar conclusion that PFA is not sensitive to arteries, nerves, esophagus, trachea and other tissues^[10-14]^. The common feature of these tissues is that they are composed of more fibrous tissues, such as elastic fibers, nerve fibers and fibrous connective tissues. Atherosclerosis is initiated by disease of the intima of the arteries. Foam cells formed by macrophages or smooth muscle cells engulfing lipids thicken the intima and thin the tunica media^[15]^. The significant difference in the content of elastic fibers between the thickened intima and the thinned media leads us to assume that PFA has the potential to treat local atherosclerosis. As expected, our results confirm that PFA 2000V has a distinct effect (apoptosis and polarization alignment) on the atherosclerotic intima, but not on normal arterial vessels (figure3). As expected, our results confirm that PFA2000V has a significant effect (apoptosis and polarization arrangement) on the atherosclerotic intima, but has no significant effect on the thinned but elastic fiber-supported media. Of course, PFA also has no ablative effect on normal arterial vessels, possibly because there is only a endothelial monolayer there.

Lipid dysmetabolism is the primary factor in the formation of atherosclerosis. Although the new view is that in many patients treated for atherosclerotic cardiovascular disease, elevations of triglyceride-rich lipoprotein (TGRL) and low-high-density lipoprotein (HDL) constitute the dominant pattern of lipid abnormalities, rather than elevations of low-density lipoprotein (LDL) cholesterol^[16]^. In animal models, apolipoprotein B, the signature component of LDL, is still routinely used to measure lipid deposition. In our results, lipid deposition was significantly reduced and was only present in the nascent media (figure4), while the inner membrane remained monolayer cell structure. This phenomenon has not been reported in either stent or balloon dilation therapy^[17-19]^. The reason for this is not yet known, but the newly born media on day 7 after PFA ablation has a very active proliferative capacity(figure5), which may indicate the presence of phagocytic cells.

The rapamycin-loaded balloon is often used to dilate the stenosis of blood vessels caused by atherosclerosis. The effect of rapamycin is generally thought to inhibit the proliferation of macrophages or smooth muscle cells and thus prevent the continued expansion of plaques^[20]^. In this study, PFA combined with rapamycin was also intended to inhibit possible cell proliferation following PFA destruction of foam cells in atherosclerosis. The inhibitory effect of rapamycin on cell proliferation was clearly observed in our study(figure5). However, this effect did not play a synergistic role with PFA, but delayed the vascular remodeling induced by PFA.

We have not seen other reports on the ablation of atherosclerosis by PFA, but from our study, the remodeling of vascular structure may be an important mechanism for the use of PFA in the treatment of atherosclerosis. After PFA ablation, blood vessels undergo a process of structural remodeling, in which the newly formed smooth muscle layer with the characteristic of polarization gradually replaces the original atherosclerotic intima (figure6). The atherosclerotic tissue that has not yet been replaced is sandwiched between the new layers of smooth muscle and the remaining media. Both PFA parameters and rapamycin can affect the rate of replacement. It suggests that optimizing PFA ablation parameters or combining lipid-lowering drugs or other drug therapy may be the future research direction if we want to obtain complete replacement of the media layer.

## 5. Conclusion

PFA is a novel ablation technique that we are trying for the first time to apply to the local treatment of atherosclerosis. We performed PFA on a rabbit atherosclerotic model and confirmed that PFA can effectively reduce lipid deposition in atherosclerotic plaques, and most importantly, promote vascular remodeling in the atherosclerotic damaged area, increase the smooth muscle media thickness of new vessels, and thus enhance the stability of blood vessel walls. Our study suggests that PFA is a promising approach for the treatment of atherosclerotic plaques. Of course, our study is still very preliminary, and future studies need to elucidate the mechanism of PFA ablation of atherosclerosis, observe the long-term effects, and optimize the ablation regimen.

### Disclosures

The authors Su Siying and Liu Shangzhong are employed by Tianjin Intelligent Health Medical Co., Ltd, and the PFA equipment in this study is provided by Tianjin Intelligent Health Medical Co., Ltd. The other authors declare no conflict of interest.

## Author Contributions

Conceptualization, Yin Huijuan, Xue Zhixiao and Lu Chengzhi; methodology and investigation, Ye Xuying, Hu Jiashen, Cao Shisheng; data curation, Ye Xuying; writing—original draft preparation, Yin Huijuan; writing—review and editing, Xu Xinyu; visualization, Yin Huijuan; funding acquisition, Ye Xuying. All authors have read and agreed to the published version of the manuscript.

## Funding

The research was funded by Tianjin Health Commission (Number TJWJ2022MS017).

## Notes

### Competing Interest Statement

The authors have declared no competing interest.

